# Design and deployment of an affordable and long-lasting deep-water subsurface fish aggregation device

**DOI:** 10.1101/2021.01.23.427792

**Authors:** Eric VC Schneider, Edward J Brooks, Michael P Cortina, David M Bailey, Shaun S Killen, Travis E Van Leeuwen

## Abstract

Fish aggregation devices (FADs) are used worldwide to enhance the efficiency of various fisheries. Devices consist of a floating or subsurface component designed to exploit natural fish behavior, using species’ attraction to structure (e.g., *Sargassum* spp.) to aggregate fish and increase capture success in open ocean environments. Concerns have arisen regarding the scale and management of FAD-associated fisheries, however, the efficiency of FADs to aggregate fish also introduces the possibility for FADs to be used as conservation tools to study pelagic species ecology. Building on two successful and several failed deployments of anchored deep-water (>500 m) subsurface (10 m) FADs over three years in The Bahamas, and observations from the subsequent FAD monitoring program, the objectives of the paper are to: 1) provide details and considerations for the design, construction, and deployment of an affordable and durable deep-water subsurface FAD that can be deployed using small boats; and 2) highlight the potential for a long-lasting moored FAD to be used as a sustainable and reliable scientific platform for pelagic species research and conservation, lending specifically to several research applications. This information will be useful for assessing the impacts that FADs and other anthropogenic marine infrastructure have on wild marine species, and their efficacy for conserving pelagic fish through increased encounters for study.

## Introduction

The pelagic ocean is the largest habitat on earth by both surface area and volume, however, it is largely understudied compared to coastal or terrestrial ecosystems (Webb et al. 2010). The pelagic zone of the ocean provides important ecosystem services such as food and oxygen production, carbon cycling, and climate stabilization (Robison 2009), and is known to harbor considerable biodiversity (Angel 1993). With offshore habitats under intense fishing pressures (Dulvy et al. 2008, Verity et al. 2002) and many fish stocks reaching either fully exploited or over-exploited levels (FAO 2018, Pons et al. 2017), the need for better science, management, and enforcement of the pelagic zone and its fisheries is evident and increasing.

One aspect of fish behavior, particularly seen in pelagic species, that has substantially contributed to their harvest is their propensity to aggregate around floating structure (Castro et al. 2002, Girard et al. 2004). Over the past several decades, intentionally constructed fish aggregation devices (FADs) have become a ubiquitous tool in pelagic purse seine fisheries (Moreno et al. 2016) with more than half of all tuna landed globally caught using FADs (Miyake et al. 2010). A wide range of epi-pelagic fishes have been documented aggregating to floating structures (Castro et al. 2002) including many high-level predators such as scombrids (e.g., tunas), billfish, and pelagic sharks which are groups of fishes most at risk of over-exploitation (Baum et al. 2003; Baum and Myers 2004). It is estimated that 81,000 to 121,000 new FADs are deployed into the world’s oceans annually, with many lost within the first year after deployment (Gershman et al. 2015). The scale of FAD use poses numerous challenges to ocean conservation and fisheries management, and such a rapid increase in fishing technology (i.e., FADs instrumented with GPS trackers and ‘fish-finder’ echosounders) have outpaced management developments (Baske et al. 2012). When working to manage an industry that utilizes such powerful fishing tools as FADs, extra attention must be applied to promote the sustainability of targeted stocks and bycatch.

Recently, there has been a concerted effort to utilize instrumented FADs and to work cooperatively with fishing industries to increase the capacity for pelagic species research in this often difficult to access and vast habitat (Brehmer et al. 2019, Davies et al. 2014). However, these initiatives typically use FADs that are actively fished and free-floating which may bias ecological research towards geographical areas that are chosen by fishing fleets and potentially mask ecological phenomena due to the nature of this extractive process. Additionally, most drifting FADs are not accessible by small boats with shorter ranges and have a relatively short functional life of less than one year before degrading or washing out of range and / or ashore, limiting the feasibility of long-term biological studies or oceanographic monitoring (Lopez et al. 2017). Therefore, to address these concerns, it is important to invest in the collection of fisheries-independent data utilizing long-lasting anchored FADs to better understand the status and trends of commercially important fish stocks (Moreno et al. 2016). Depending on location, anchored FADs may allow greater accessibility to undertake monitoring work, can facilitate a wide array of instrumentation both above and below the surface of the ocean, and will allow for longer studies to occur if designed properly. Few resources exist on the design, construction, and deployment of anchored FADs for study, despite their widespread use in the fishing industry. Existing studies and manuals typically describe FADs deployed from large boats or using materials that are either expensive or difficult to purchase and ship to remote locations (see examples in Table 1).

**Table 1.** Selected examples from a search of studies using anchored FADs that include information on materials or deployment, that are subsurface (or ‘midwater’), or that were deployed in depths ≥ 500 m and therefore comparable to our proposed design. ‘Buoy Depth’ of 0 represents a surface FAD. Substantial and replicable information must be included on materials used or deployment processes to qualify as ‘Yes’.

| FAD Type | Bottom Depth (m) | Buoy Depth (m) | Materials Info | Deployment Info | Location | Reference |
| --- | --- | --- | --- | --- | --- | --- |
| Surface and subsurface | 300-2000 | 0-? | Yes | Yes | Pacific Islands | Chapman et al., 2005 |
| Surface | <200 | 0 | Yes | Yes | Timor-Leste | Mills & Tilley, 2019 |
| Surface | 100-3000 | 0 | Yes | Yes | Caribbean | Gervain et al., 2015 |
| Surface and subsurface | 1000-2000 | 50-100 | Yes | No | Japan | Sokimi, 2006 |
| Surface and subsurface | 146-2761 | 0-18 | Yes | No | Hawaii | Higashi, 1994 |
| Surface | <5000 | 0 | Yes | No | Pacific Islands | Itano et al., 2004 |
| Subsurface | 415 | 50 | No | No | Taiwan | Weng et al., 2013 |
| Surface and subsurface | 50-2500 | ? | No | No | Pacific Islands | Taquet et al., 2011 |
| Surface and subsurface | 300-700 | 0-? | No | No | Pacific Islands | Bell et al., 2015 |
| Surface | 260-600 | 0 | No | No | Puerto Rico | Merten et al., 2018 |
| Surface | 50-500 | 0 | No | No | Canary Islands | Castro et al., 1999 |
| Surface | 1000-2200 | 0 | No | No | American Samoa | Buckley & Miller, 1994 |
| Surface | 2000-2500 | 0 | No | No | Martinique | Doray et al., 2007 |

Here we detail the design, construction, deployment, and utility of an anchored deep-water, subsurface FAD that is durable and long-lasting, easily reproducible, and specifically intended to facilitate fish ecology and marine conservation research. The durability and moored nature of the design makes the FAD less prone to loss or damage, and the low cost of the FAD allows for the possibility of several to be deployed using small boats and facilitate much needed replication and manipulation in experimental designs, a component often lacking in this area of study. While the subsurface aspect of these FADs was designed to reduce surface-associated damage and tampering, and to facilitate specific research objectives, any potential trade-offs between constructing a subsurface FAD and fish attraction / aggregation are also a point of interest and are detailed below.

## Methods

### FAD Design

The main design objectives were to create a long-lasting subsurface anchored FAD without any surface markers, deep enough to avoid surge-associated damage or navigational hazards, while still shallow enough to be accessible to divers and avoid pressure-related damage to the steel buoys. Additionally, the FAD needed to easily facilitate equipment mounting to act as a stable platform for various fish ecology and marine conservation investigations.

The FADs used in our study consisted of a concrete anchor block (122 cm L x 122 cm W x 84 cm H = 1.25 m^3^; ∼2900 kg weight on land, ∼1620 kg weight in water), a vertical mooring line (600 m of 5,817 kg minimum tensile strength one inch polypropylene) and two tethered subsurface steel buoys (surplus naval buoys, 71 cm diameter, 54 kg weight / 181 kg buoyancy each; Fig. 1). A depth of 10-15 m subsurface was decided to be an acceptable target range for the floats, although subsurface FADs are not commonly deployed at bottom depths > 600 m (study site depth) which is novel here (Chapman et al. 2005). At the time of writing, two of these FADs have remained in place for over 3 years (December 2017 – May 2021) and have withstood wind and surge from multiple passing hurricanes (<100 kph winds). The FADs were visited weekly for the first 2.5 years, and then monthly thereafter.

**Figure 1.**
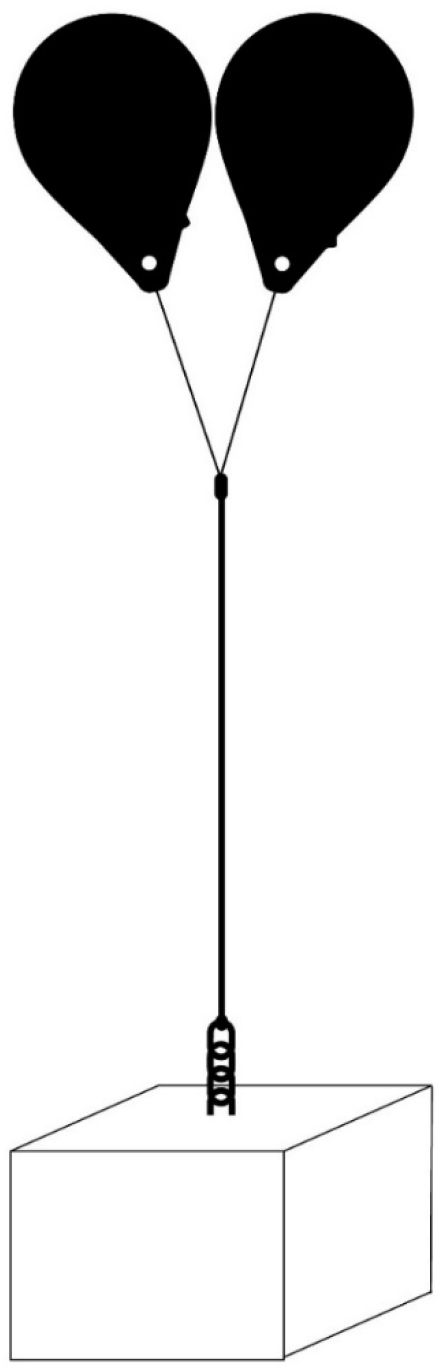
Schematic of the subsurface fish aggregation device (FAD) currently being used at The Cape Eleuthera Institute, The Bahamas. FAD design consists of a concrete anchor block, mooring line, and two steel buoys. Steel buoys are moored 10m below the sea surface to prevent detection by fishers and to create tension and verticality in the mooring line for gear deployment. Diagram not to scale.

Existing designs found in scientific papers and technical reports did not conform to all our objectives, therefore a new and unique design was utilized here. For example, Weng et al. (2013) used a subsurface FAD at a bottom depth of 415 m, however, the buoy depth of 50 m was inaccessible to divers. Further, details of an array of subsurface FADs around Okinawa, Japan indicate subsurface FADs at bottom depths up to 2000 m. However, the materials and deployment were costly, and structure depth ranged from 20 m to 100 m subsurface which minimizes survey time available for divers or renders it inaccessible (Sokimi 2006). The closest example to those used here is a subsurface FAD array in Hawaii described by Higashi (1994) with a bottom depth range of 366 m to 549 m and a buoy depth of 18 m to 21 m, but little detail describing the construction and deployment are available. From the limited information available, it suggests a large naval vessel was used for deployment and that the FAD construction used galvanized steel cable, suggesting neither a simple nor cheap construction and deployment process.

### Construction

Steel reinforcing lattice (#4 rebar) was laid as the concrete was being placed, and a stainless-steel round stock bail (Fig. 2) was incorporated beneath the last layer of lattice so that the top of the bail protruded from the center of the block to aid in mooring line attachment. A shackle (7/8” bolt through) was used to attach 4 meters of 3/4” long-link chain to the bail anchor point between the block and mooring line. The chain was used to prevent chaffing of the anchor line against the anchor block in the event of converting the structure to a surface FAD. However, this was determined to be unnecessary for subsurface orientation because the tension in the mooring line, by default, prevents the anchor line from contacting the anchor block. This was followed by a 7/8” bolt through shackle, a 7/8” eye by eye swivel, another 7/8” shackle and finally a size 4 rope connector (Samson Nylite, Ferndale, Washington, USA; Fig. 3).

**Figure 2.**
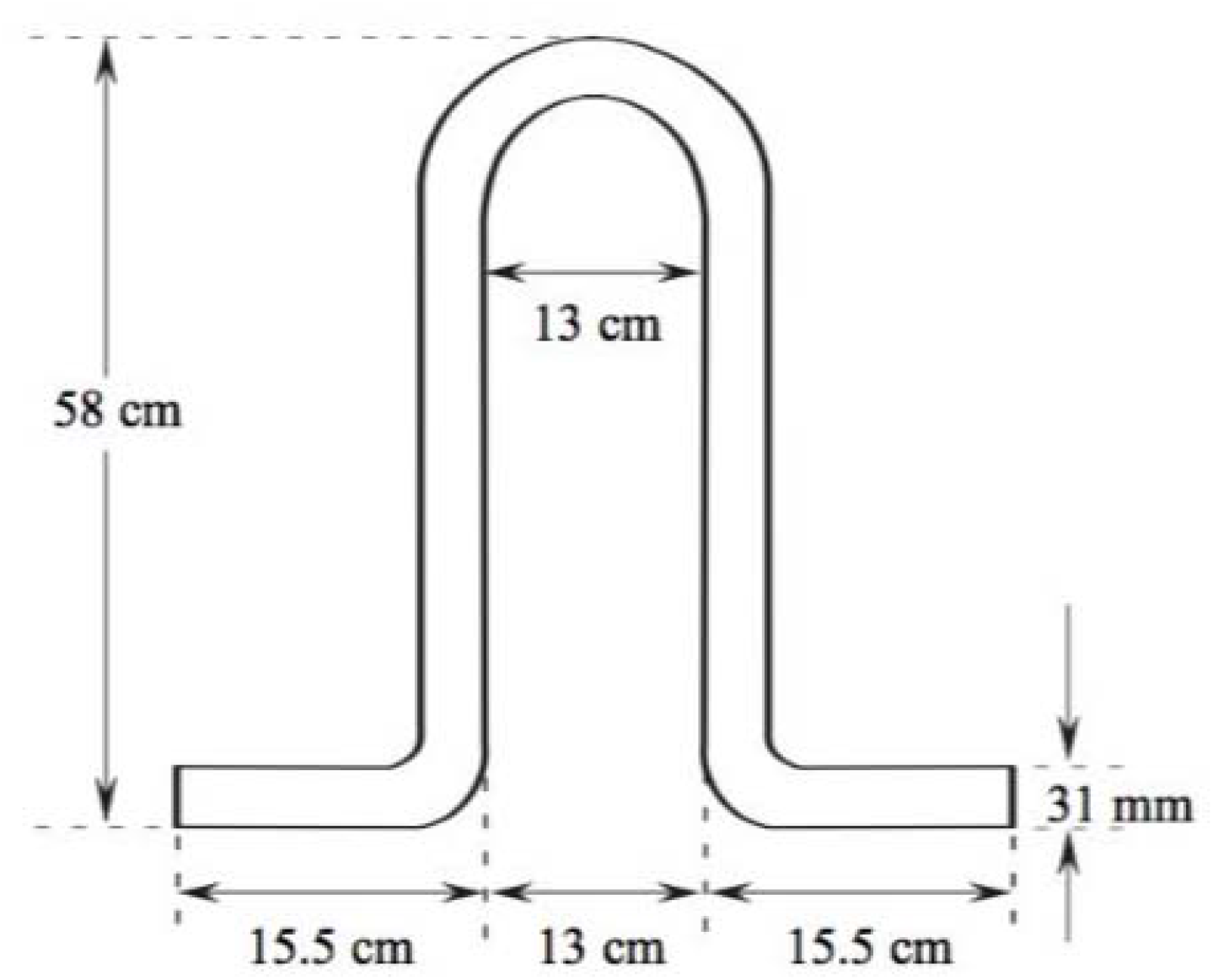
Stainless steel round stock bail that was incorporated into the top of the concrete anchor block to serve as the attachment point between the block and metal chain, which was the beginning of the mooring line.

**Figure 3.**
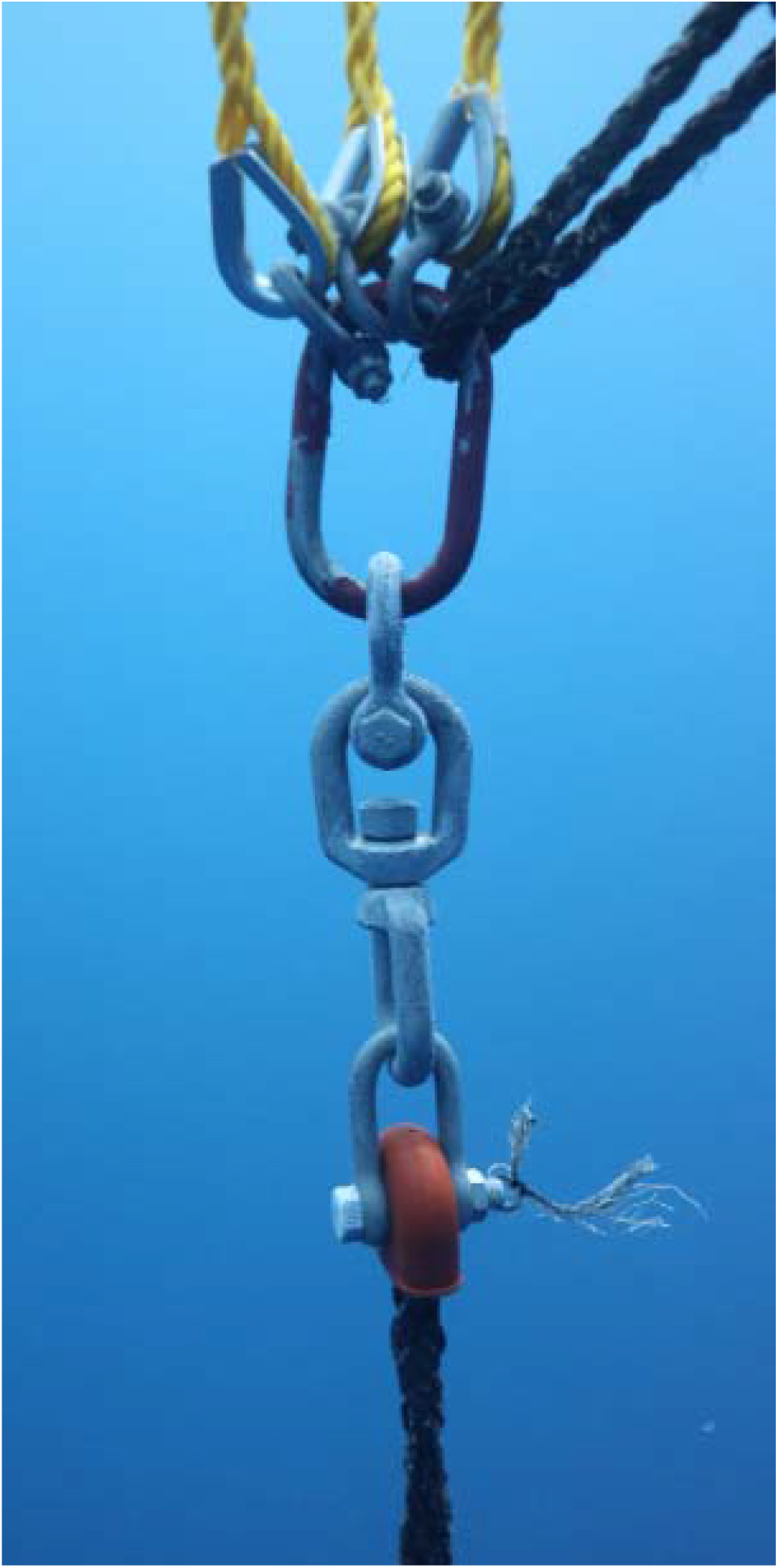
Top end of the mooring line showing an eye splice to a rope connector, swivel, bolt through shackle, and 7/8” master link to which the buoys (and a safety line during deployment) were attached.

Eight-strand 1” polypropylene line (5,817 kg minimum tensile strength) was used as the mooring line and attached to rope connectors via an eye-splice. The rope connector prevented the eye splice from chaffing against the metal rigging. End-to-end splices were used any time the line needed to be extended. The same series of hardware was repeated at the end of the line (rope connector, swivel, and shackle). However, this series was then followed by a 7/8” master link (Fig. 3).

The floating portion of the FAD structure was comprised of two round 71 cm diameter steel buoys (54 kg weight, 181 kg buoyancy each) tethered to the master link using 2 meters of 1/2” three-strand polypropylene line that was eye-spliced through a rope thimble. This is similar in size and surface area to the flotation component of drifting FADs used in some commercial tuna fisheries (personal observation). However, FADs used in commercial fisheries often incorporate subsurface netting, palm fronds, synthetic streamers, or other structure below the surface. These additions were not included in this design to avoid animal entanglement (Filmalter et al. 2013) and to maintain clearance along the mooring line for equipment deployment (hydrophones, acoustic receivers and oceanographic monitoring equipment) and retrieval during future stages of the research program.

### Deployment

Several location parameters were taken into consideration when selecting locations for the subsurface FADs, and the Exuma Sound (near the Cape Eleuthera Institute and base of research operations) was ideally suited for this. First, a deep-water drop off was located near-shore and accessible from the research station. Second, the area is a known migration route for pelagic fishes. Although the bathymetry had not been accurately described, several known depth points from previous deep-water research were used to select suitable locations and to predetermine mooring line lengths.

Following the construction of the individual components on land, the anchor was transported into shallow water at a nearby marina using a crane truck. Although in this case a crane truck was used, the block could be constructed on a platform at the edge of the water and deployed using rollers or a winch for simplicity. Once the anchor was submerged, three lift bags (SP2000, Subsalve, North Kingstown, Rhode Island, USA) were attached to the anchor bail using a release under load mechanism (Sea Catch TR7, MacMillan Design, Gig Harbor, Washington, USA). A safety line was attached between the block and lift bag to prevent premature deployment. Lift bags were inflated with compressed air and the anchor raised off the bottom for towing behind a small boat. The tow line was attached above the release mechanism so that if the anchor block dropped unexpectedly the weight of the anchor would not damage the boat. Once at the deployment location, a second small vessel slowly released the mooring line overboard and onto the surface of the water down current and away from the anchor attachment point. A polyethylene ball float was attached to the free end of the mooring line to aid in visualization of the rope during and after deployment. Following the deployment of the mooring line from the vessel, snorkelers attached the anchor chain to the bail using a shackle, removed the safety line between the block and the lift bags, and released the load-bearing mechanism using a nylon rip cord thus dropping the block (Fig. 4). The location of the drop was positioned to be 1/3 of the depth up-current of the targeted FAD location to account for drag on the line pulling the block in the down-current direction.

**Figure 4.**
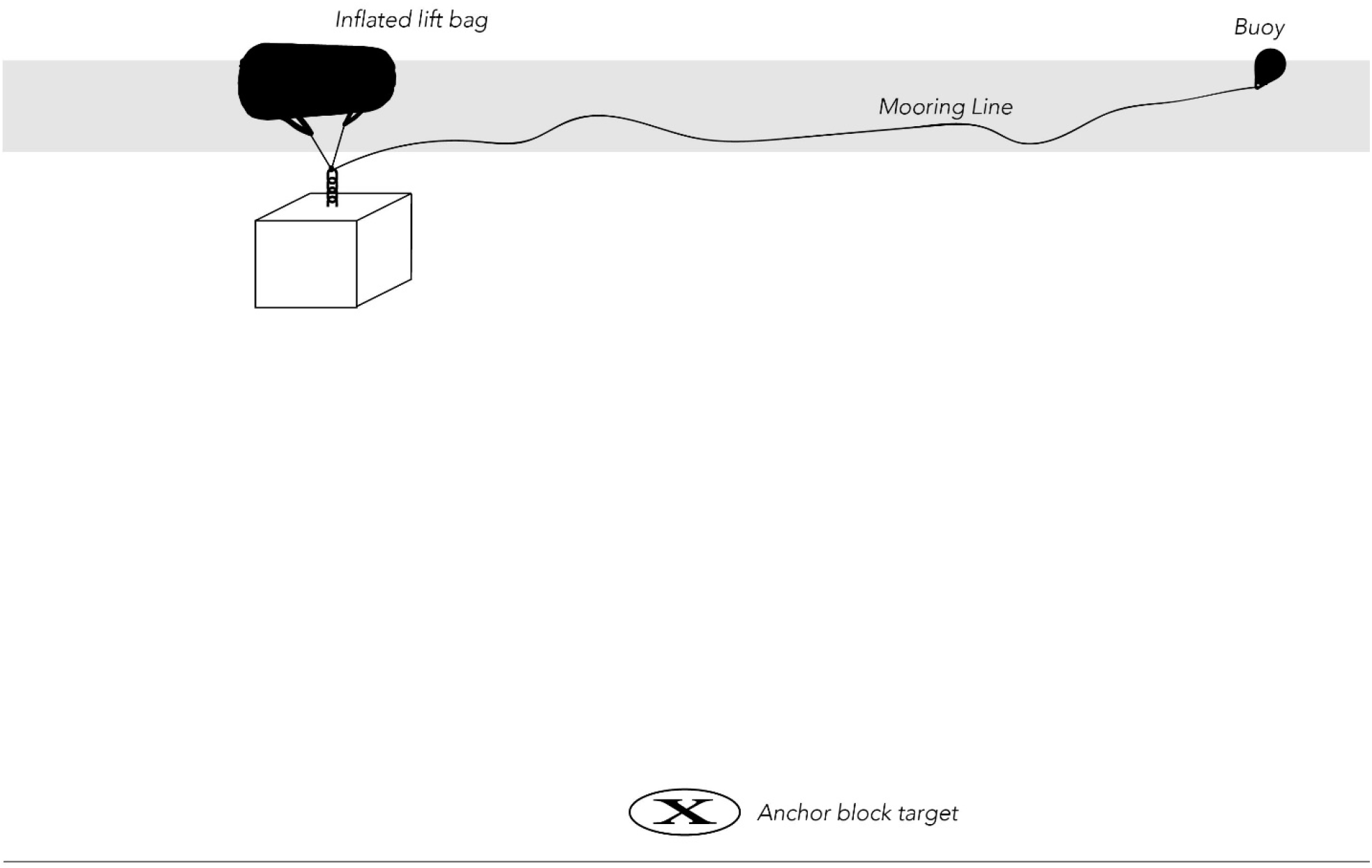
Initial stage of the FAD deployment process. The concrete anchor block is suspended by lift bags and attached to the mooring line that has been deployed overboard onto the sea surface down-current from the anchor. The anchor is positioned 1/3 the length of the mooring line up-current of the targeted resting location.

Following the anchor drop, the excess mooring line on the surface was recovered. The mooring line was then elongated using lift bags attached at depth to simulate the ultimate tension on the line from the FAD buoys. To do so, divers on SCUBA attached a lift bag to the mooring line using a Prusik hitch at 25 m depth and filled the lift bag with compressed air (Fig. 5). Following the lift bag’s ascent to the surface, this process was continued until the line was under approximately 600 kg of tension, evidenced by the 900 kg lift bag filled to approximately 66% of total volume, and positioned at a static depth of 10 m without further elongation. At least 24 hours were allowed for a complete tidal cycle, and to allow the mooring line to undergo phase one creep (stretching) to its ultimate length under load. If this time period resulted in reduced tension, or if the lift bags reached the surface, the mooring line elongation process was repeated. When the line was determined to have undergone all elongation, an eye-splice was used to attach a rope protector to the mooring line just above the lift bag. A shackle, master link, and a 15 m safety line with a fully inflated SP2000 lift bag were attached to the trailing end of the mooring line at the surface to further ensure the mooring line did not retract. Two individually rigged steel buoys were spliced onto the master link. After the attachment was complete, the lift bag under tension was released allowing the recoil of the mooring line to submerge the buoys to a depth of 10 to 15 m, at which point the inflated surface lift bag prevented any possible further descent. A small polyethylene buoy was finally attached to the master link and filled with compressed air as needed to fine tune buoyancy to the targeted 10 m depth resting point. Lastly, a safety line (3/8” Spectra 12-strand braided line, 6,305 kg minimum tensile strength) was tied to the master link, run through each eye-attachment point on the two buoys and tied back to the master link. In the case of eye-ring failure or a buoy tether being severed, this line provides a cut-resistant back-up to avoid the loss of a buoy or the entire FAD.

**Figure 5.**
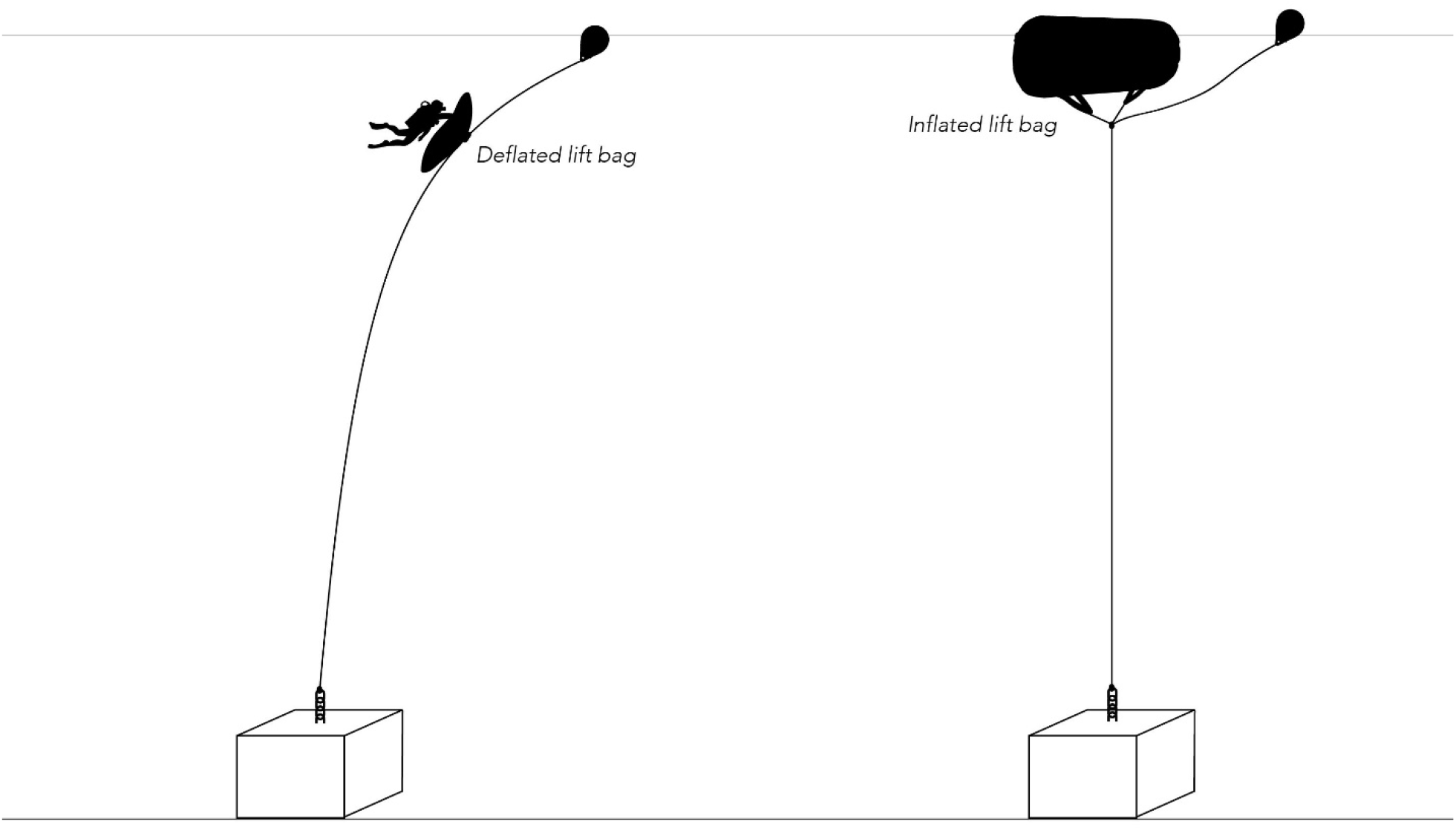
Schematic showing the process of removing slack from the mooring line following deployment. A diver attaches a deflated lift bag to the slack mooring line using a Prusik hitch and slowly inflates the bag. This process is repeated until the desired tension on the mooring line has been reached and the steel buoys are then attached.

### Removal Potential

While the durability and moored nature of the FAD design presented here results in a robust and maintainable platform for longer time scales, these FADs are removable using the same equipment and process needed for deployment and users can therefore avoid contributing marine debris to the ocean following the conclusion of research activities. Although this would involve some effort, divers repeatedly deploying lift bags down the mooring line will slowly raise the anchor block which can then be towed to shallow water and allow for retrieval of the entire FAD.

## Results and Discussion

### Design Considerations

Materials and operations were all considered and selected to not only meet the project objectives, but to be accessible by a wide array of potential users including those at remote field stations or research groups with limited funding and resources. It has been shown that price is often a limiting factor during FAD creation and installation in remote island locations, and although this is typically documented in the scope of bolstering fishing communities (Bell et al. 2015), financial restraints will similarly apply to research groups. During the development and expansion of the Pacific FAD fishery, 2000 to 3000 USD was targeted for a reasonable total cost for a deep-water FAD intended to last approximately 2 years (Chapman et al. 2005), so the 5000 USD total cost per FAD in this project was deemed appropriate when designing for a durable longer-lasting structure. Longevity is also a high concern for surface FADs, with wave/weather damage or vandalism frequently leading to loss of the FAD in less than two years (Chapman et al. 2005, Tilley et al. 2019). Inspections in June 2019 (one and a half years after deployment), using deep sea submersible surveys in the area, revealed that all parts of the FAD design inaccessible to divers remain in good condition. The concrete anchor block, steel connections (shackles, chain, swivels, etc.), polypropylene mooring line and steel buoys were all considered to be affordable and possible to source and ship to a remote location and are standard options for offshore FADs (Chapman et al. 2005). Recently, there has been considerable effort to construct FADs from biodegradable materials to decrease marine debris and the potential negative impacts on wildlife such as entanglement (Moreno et al. 2018). Increasing longevity through careful design and robust synthetic materials was pursued during this project, although this could easily be adapted for a shorter-lived but biodegradable version. Additionally, the FADs were deployed using only SCUBA divers, a crane truck (or equivalent for pushing the anchor into the water), lift bags, and two 8 m long inboard panga vessels. One possible price reduction was tested by using three A-6 sized polyethylene buoys (Polyform A-6) instead of steel buoys, however this was quickly proven to not work. Flexible buoys expand or contract with minimal changes in water depth associated with the mooring line stretching or contracting, which in turn changes the buoyancy and prevents a stable target depth. Additionally, several flexible buoys showed marks consistent with teeth punctures by predatory fishes. Therefore, steel buoys soon replaced the flexible buoys after deployment of the first FAD and are highly recommended. Alternative mooring line materials were also considered, and materials such as Spectra or Dyneema are cut-resistant and would dramatically reduce phase 1 creep (stretching), making buoy placement at a target depth easier, however, these options are considerably more expensive and were avoided for this reason.

The subsurface aspect of the FAD design was chosen for several reasons related to the objectives of the project. First, 10 m depth was found to greatly minimize movement of the structure by surge or during windy weather, and would prevent any potential boat strikes, adding to the longevity of the infrastructure. Additionally, 10 m is an easily accessible and safe depth for both SCUBA and freedivers to work or deploy/retrieve equipment and does not pose any serious pressure-related stress to the buoys or equipment (at 2 atmospheres).

## Research Applications and Conclusions

The subsurface anchored design of the FADs used in this study has proven to be a stable and diverse scientific platform for more than three years. Many of the epipelagic fish species that occur in the region have been documented at the FADs, ranging in trophic level and size (Table 2). These anchored FADs allow the fish community to be continually monitored over long temporal scales and can facilitate short and long-term experimental studies through increased accessibility that would not be possible when using conventional offshore drifting FADs. It is possible that these long-term stationary fish censuses could be representative of actual population trends, and these data would therefore be useful to fisheries managers. Additionally, Dagorn et al. (2010) previously argued that anchored FADs are acceptable and useful proxies for drifting FADs to address research questions such as the ecological trap hypothesis, and that they pose accessibility advantages while maintaining contextual similarities to their drifting counterparts. A variety of methods that have been recently performed on surface FADs and would be well-suited to this FAD design, many of which were proposed by Moreno et al. (2016) as research priorities, are detailed in Table 3.

**Table 2.** Species documented on these subsurface FADs in the Exuma Sound during the course of a 2.5 year camera survey (in preparation for publication elsewhere), separated into resident intranatant versus ephemeral circumnatant species, compared to other epipelagic fishes documented in the Exuma Sound (personal communication: Z. Zuckerman, Cape Eleuthera Institute) but remain absent from our subsurface FAD surveys. An asterisk (*) denotes species present within first 6 months after FAD deployment.

| Present on subsurface FAD surveys |  | Absent from subsurface FAD surveys |
| --- | --- | --- |
| Intranatant | Circumnatant | Recorded in Exuma Sound |
| *Aluterus Monoceros* * | *Acanthocybium solandri* * | *Carcharhinus longimanus* |
| *Balistes capriscus* * | *Carcharhinus falciformis* * | *Istiophorus albicans* |
| *Cantherhines* spp.* | *Carcharhinus obscurus* | *Kajikia albidus* |
| *Canthidermis sufflamen* * | *Cheilopogon melanurus* | *Makaira nigricans* |
| *Carangidae* spp.* | *Coryphaena hippurus* * | *Scomberomorus cavalla* |
| *Caranx latus* * | *Elagatis bipinnulata* * |  |
| *Caranx ruber* * | *Galeocerdo cuvier* * |  |
| *Decapterus* spp.* | *Hemiramphus* spp. |  |
| *Decapterus macarellus* * | *Sarda sarda* |  |
| *Peprilus triacanthus* | *Sphyraena barracuda* * |  |
| *Seriola rivoliana* * | *Sphyrna mokorran* |  |
|  | *Thunnus albacares* |  |
|  | *Thunnus* spp. |  |

**Table 3.**
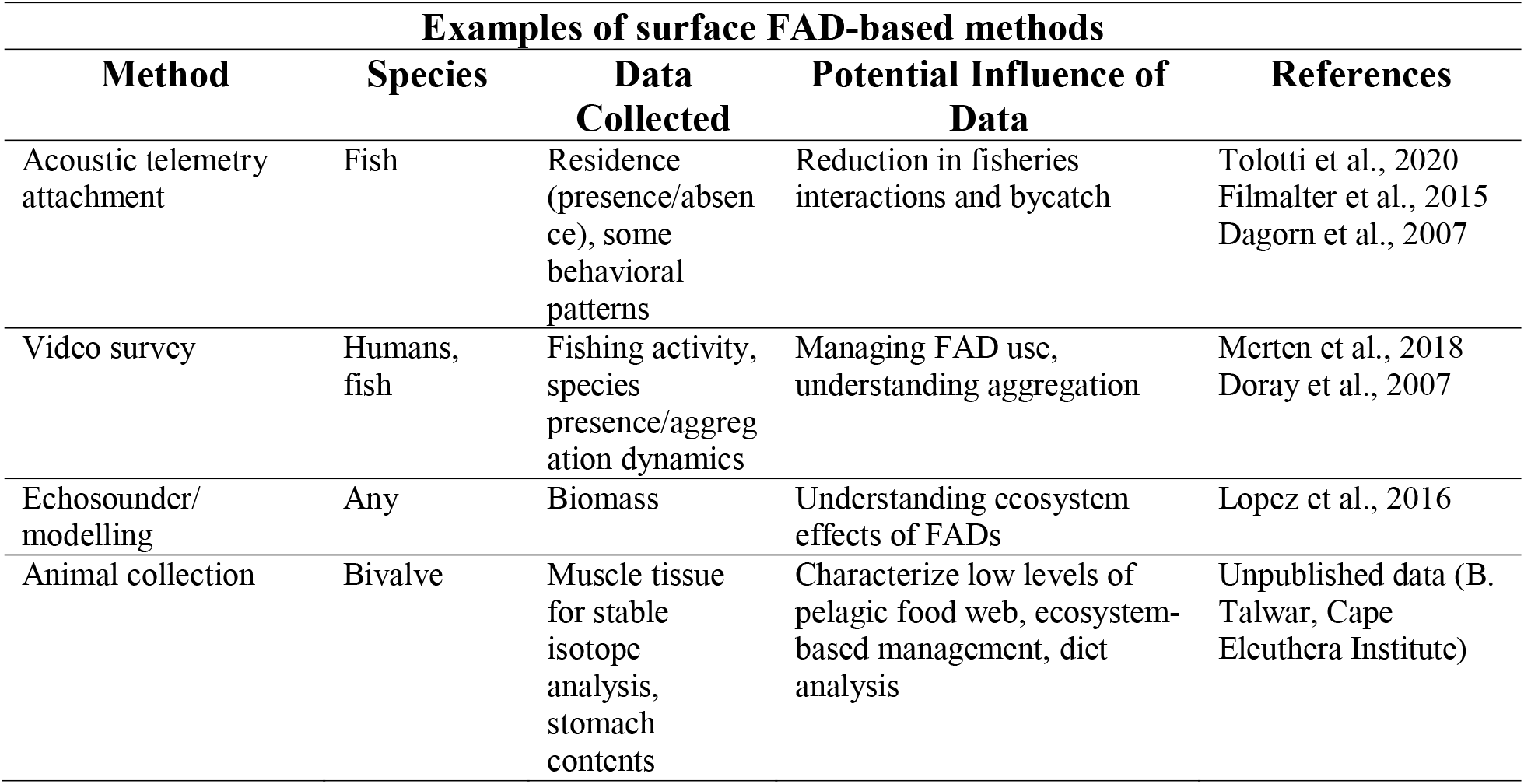
Recently utilized methods for FAD-based research that lend well to a stable, subsurface platform.

The subsurface aspect of the FADs used in this study, combined with the buoyancy of the buoys used (362 kg of lift total), resulted in a taut mooring line. Therefore, this design presents several unique opportunities for research activities. First, equipment can be shuttled up and down the mooring line with a simple rigging system, enabling fixed depth/location deployment of various instruments. This would otherwise be difficult without a taut mooring line. Additionally, keeping the structure underneath the surface and away from any surge or waves results in a nearly silent structure, presenting opportunities for investigation into fish sensory biology in the open ocean, and for better hydroacoustic data collection (e.g., hydrophone deployment for cetacean surveys). Finally, this tension ensured that the FAD buoys remained at the known GPS location and did not sway with tidal or current flow. Although this study area does not experience significant currents, locations with strong tidal flow should consider the impact of horizontal forces on the FAD and mooring line.

Strategically designed FADs that are not open access can act as useful research platforms to develop new monitoring approaches while maintaining the integrity of the study population and enhance our understanding of how anthropogenic activity is affecting marine biodiversity. For example, many animals that utilize pelagic FADs are fish species known to undergo long migrations (Hallier and Gaertner 2008) which can be difficult to study. Whether following seasonal changes in food abundance, thermal windows, or breeding opportunities (Alerstam et al. 2003), this migratory behavior most likely exposes them to various fisheries pressures and potential overexploitation. Knowledge of how migratory animals respond to variable conditions experienced during migration is a central component to understanding long-distance movement patterns and their management. If data are collected consistently, this information can help estimate population size, increase understanding of demographic variables needed in the development of population viability models, reveal how wild fish species are impacted by anthropogenically altered habitats, and can potentially be used for novel conservation applications. These include the construction of scientific platforms (such as deep-water subsurface FADs) along known migration routes to aid in the study of elusive migratory animals, or the ability to alter movements of migratory animals through protected seascapes by enhancing habitat preferences in these areas to minimize harvest. By utilizing FAD-based equipment such as video cameras or acoustic telemetry receivers, information on behavior during migration can be collected and used in fisheries conservation.

Pelagic animals are inherently difficult to study due to the expanse of their habitat, life history, and behavior, resulting in a comparatively weak understanding of pelagic species ecology and biology (Block et al. 2003). Therefore, in response to the recent call for developing methods to collect fisheries-independent data to be used in management and stock assessments (Moreno et al. 2016), a network of economical, instrumented, research-oriented subsurface FADs such as those proposed here could provide substantial ecological and fisheries data that is desperately needed to effectively conserve pelagic ecosystems and their biodiversity.

## Authors’ Contributions

TV, ES conceptualized the manuscript; ES, TV, MC wrote the manuscript and all authors contributed substantially to revisions and accept responsibility for this work.

## Acknowledgments

This is contribution #1 of the Exuma Sound Ecosystem Research Project. We would like to thank The Moore Charitable Foundation / Moore Bahamas Foundation for generously funding this work. S.S. Killen was supported by a NERC Standard Grant NE/T008334/1. We extend many thanks to the staff, interns, students, and visiting researchers of the Cape Eleuthera Institute and The Island School for their extensive assistance including M. Israel, D. Huber, P. Osborn, C. Hsia, D. Orrell, E. Good, G. Sayles, D. Grady, and W. Barnes, in addition to the students of The Island School FAD research classes. We thank L. Madden for assistance in creation of all graphics included in the figures. The authors have no competing interests.

